# Development of deaminase-free T-to-S base editor and C-to-G base editor by engineered human uracil DNA glycosylase

**DOI:** 10.1101/2024.01.01.573809

**Authors:** Huawei Tong, Haoqiang Wang, Nana Liu, Guoling Li, Yingsi Zhou, Danni Wu, Yun Li, Ming Jin, Xuchen Wang, Hengbin Li, Yinghui Wei, Yuan Yuan, Linyu Shi, Xuan Yao, Hui Yang

## Abstract

DNA base editors could enable direct editing of adenine (A), cytosine (C), or guanine (G), but there is no base editor for direct thymine (T) editing currently. Here, by fusing Cas9 nickase (nCas9) with engineered human uracil DNA glycosylase (UNG) variants, we developed a deaminase-free glycosylase-based thymine base editor (gTBE) with the ability of direct T editing. By several rounds of UNG mutagenesis via rational screening, we demonstrated that gTBE with engineered UNG variants could achieve T editing efficiency by up to 81.5%. Furthermore, the gTBE exhibited high T-to-S (i.e., T-to-C or T-to-G) conversion ratio with up to 0.97 in cultured human cells. Using similar strategy, we developed a deaminase-free cytosine base editor (gCBE) facilitating specifically direct C editing by engineered UNG with mutations different from gTBE. Thus, we provide two novel base editors, gTBE and gCBE, with corresponding engineered UNG variants, broadening the targeting scope of base editors.

---

Base editors enable single-nucleotide edits with high precision and efficiency, providing powerful tools for the fields of life science and medicine^1, 2^. Two categories of DNA base editors, deaminase-based base editor (dBE) and deaminase-free glycosylase-based base editor (gBE), have been developed to date^3^. The dBEs, including adenine base editor (ABE)^4^, cytosine base editor (CBE)^5^, and double-stranded DNA deaminase toxin A (DddA)-derived cytosine base editor (DdCBE)^6, 7^ as well as their derivatives (such as A&C-BEmax^8^, AYBE^9^, AXBE^10^ and CGBEs^11–15^), perform base editing using single-stranded or double-stranded DNA deaminase enzymes (e.g., evolved tRNA adenosine deaminase TadA, AID/APOBEC-like cytidine deaminase or DddA variants). All the base conversions by dBEs require the deamination of A or C as the first key step. Recently, we have developed a gBE enabling direct G editing (i.e., deaminase-free glycosylase-based guanine base editor, gGBE)^3^, based on engineered human N-methylpurine DNA glycosylase (MPG; also known as alkyladenine DNA glycosylase, AAG). So far, dBEs and gGBE could enable editing of adenine (A), cytosine (C), or guanine (G), but no base editor for thymine (T) editing is available now. Base conversion by deamination is impossible for T (due to the absence of amine), making the development of thymine base editor still challenged.

Here, we developed a deaminase-free glycosylase-based thymine base editor (gTBE) as well as a deaminase-free glycosylase-based cytosine base editor (gCBE), to achieve orthogonal base editing, that is gTBE for direct T editing and gCBE for direct C editing, respectively. After several rounds of mutagenesis of the uracil DNA glycosylase (UNG, or UDG) moieties, we obtained marked enhancement of editing activity for T editing and C editing, as compared with that obtained by wild-type (WT) UNG variant. We characterized the editing profile of gTBE and gCBE by targeting dozens of endogenous genomic loci in cultured mammalian cells as well as mouse embryos, demonstrating their high base editing efficiency.

## Results

### Development of orthogonal base editors based on engineered glycosylases

Encouraged by the development of gGBE in our previous study^3^, we attempted to develop thymine and cytosine base editor using the deaminase-free glycosylase-based strategy. Since the three pyrimidine bases (i.e., T, C, and U) are structurally similar, we speculated that excision of canonical T or C could be achieved by engineering certain uracil DNA glycosylase, thus triggering the BER pathway and facilitating direct T editing or C editing following the one-step generation of AP sites (Fig. 1a,b). The human UNG has been engineered to excise normal T and C in DNA using mutants Y147A and N204D (numbered from UNG1) in *Escherichia coli*, respectively^16^. Alternative splicing as well as transcription from two distinct start sites leads to two different UNG isoforms, the mitochondrial UNG1 (304 amino acids, aa) and the nuclear UNG2 (313 aa), each possessing unique N-termini that mediate translocation to the mitochondria and the nucleus, respectively^17^ (Supplementary Fig. 1). To edit the nuclear DNA, we generated two prototype gBE versions, a deaminase-free glycosylase-based thymine base editor (gTBE) and a deaminase-free glycosylase-based cytosine base editor (gCBE), by fusing the two corresponding variants of nuclear UNG2 (Y156A and N213D, equivalent to Y147A and N204D of UNG1, respectively) at the C-terminus of Cas9 D10A nickase (nCas9), respectively (Fig. 1a,c). We developed two similar intron-split EGFP reporter systems as reported previously^9^, T-to-G reporter and C-to-G reporter, to evaluate the editing activity of gTBE and gCBE, respectively (Supplementary Fig. 2a). In these reporters, the AG-to-AT or AG-to-AC inactive splicing acceptor (SA) could only be remediated with T-to-G or C-to-G conversion, thus leading to correct splicing of EGFP-coding sequence and EGFP activation (Supplementary Fig. 2b). After co-transfecting gBE vectors with the T-to-G or C-to-G reporter vector containing the corresponding single-guide RNA (sgRNA) that targets the mis-splicing mutations, we found that gTBE with UNG2-Y156A (hereafter referred to as gTBEv0.1) showed slight T-to-G conversion activity, and gCBE with UNG2-N213D (hereafter referred to as gCBEv0.1) showed slight C-to-G conversion activity (Fig. 1c-e).

**Fig. 1.**
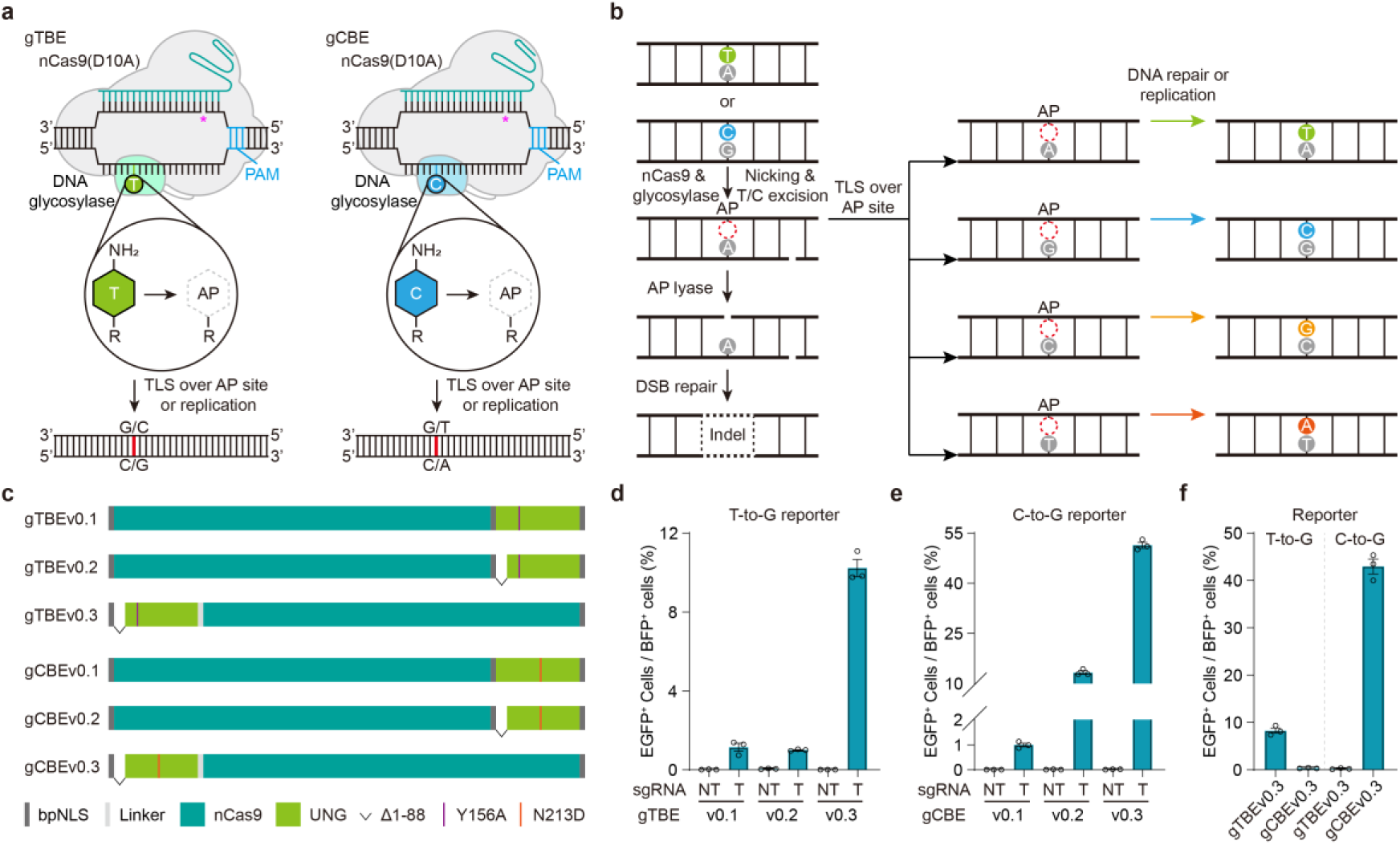
Design and mechanisms of two orthogonal glycosylase-based base editors. **a**, Prototype versions of a deaminase-free glycosylase-based thymine base editor (gTBE) and a deaminase-free glycosylase-based cytosine base editor (gCBE). PAM, Protospacer adjacent motif. AP, apurinic/apyrimidinic sites. Star (*) in magenta indicates the nick generated by nCas9. **b**, Schematic diagram of potential pathway for T (or C) editing and outcomes. A glycosylase variant is designed to remove normal T or C, an nCas9-sgRNA complex creates an R-loop at the target site and nicks the non-edited strand, then the AP site generated is repaired by translesion synthesis (TLS) and/or DNA replication, leading to T or C editing. DSB, double-strand break. indel, insertion and deletion. **c**, Schematic of various gTBE and gCBE candidate architectures. Note that Y156A and N213D of UNG2 are equivalent to Y147A and N204D of UNG1, respectively. **d**, Percentage of EGFP^+^ cells for T editing activity evaluation of different gTBE variants using T-to-G reporter (*n* = 3). NT, non-target sgRNA. T: target sgRNA. **e**, Percentage of EGFP^+^ cells for C editing activity evaluation of different gCBE variants using C-to-G reporter (*n* = 3). NT, non-target sgRNA. T: target sgRNA. **f**, the orthogonality of gTBE and gCBE for base editing evaluated using two different reporters (*n* = 3). All values are presented as mean ±s.e.m.

Given the disordered N-terminal domain (NTD) of UNG contains protein binding motifs and sites for post-translational modifications^18^, which might constrain targeted excision activity of the glycosylase domain in ssDNA^19, 20^, we constructed UNG-NTD-truncated gTBE and gCBE versions (Fig. 1c) to eliminate undesired protein-protein interactions^20–22^. The gCBEv0.2 with UNG2Δ88-N213D fused at the C-terminus increased the C-to-G conversion activity compared with gCBEv0.1, while gTBEv0.2 with UNG2Δ88-Y156A fused at the C-terminus exhibited comparable T-to-G conversion activity with gTBEv0.1 (Fig. 1c-e). Moreover, the gTBEv0.3 with UNG2Δ88-Y156A and gCBEv0.3 with UNG2Δ88-N213D fused at the N-terminus showed much higher editing activity than those at the C-terminus (10.2% vs. 1.0% and 51.4% vs. 13.3%, Fig. 1c-e), a 10- and 3.9-fold enhancement in the editing efficiency, respectively. No editing activity was found for all the above-mentioned versions of gTBE and gCBE together with the non-targeting sgRNA (Fig. 1d,e). In addition, gTBEv0.3 exhibited the highest T-to-G editing activity among various UNG-NTD-truncated versions of gTBE (Supplementary Fig. 3).

Furthermore, we examined the orthogonality of gTBE and gCBE for base editing. Although engineered from the same original glycosylase UNG, no C editing activity was found for gTBEv0.3 and no T editing activity was found for gCBEv0.3 (Fig. 1f). Thus, we developed two orthogonal base editors, gTBE for direct T editing and gCBE for C editing.

### Evolution of gTBE with enhanced editing activity

To further increase the T-to-G activity of gTBEv0.3, we attempted to perform rational mutagenesis for engineering the UNG moiety, using the T-to-G reporter to evaluate the editing activity (Fig. 2a). Based on structural and functional analysis, WT UNG contains five conserved motifs required for efficient glycosylase activity: the catalytic water-activating loop, the proline-rich loop, the uracil-binding motif, the glycine-serine motif and the leucine loop^23–25^ (Supplementary Fig. 1b). Since Y156 in the catalytic water-activating loop and N213 in the uracil-binding motif are critical for activity switch from U excision to T or C excision, we firstly selected sequential and spatial neighbors of these two residues and examined their roles in the regulation of base excision activity (Fig. 2a,b). We conducted alanine-scanning mutagenesis by replacing all non-alanine with alanine (X>A) and alanine with valine (A>V) to cover all the residues in the regions of I150-L179 and L210-T217. Interestingly, we obtained a variant gTBEv1.1 (v0.3 with A214V) largely elevating the T-to-G conversion activity by 2.68-fold (Supplementary Fig. 4a). With further site-saturation mutagenesis focused on the residue at position 214, we generated gTBEv1.2 (v0.3 with A214T) with elevated editing efficiency by 1.06-fold in comparison with the T editing activity of gTBEv1.1 (Supplementary Fig. 4b).

**Fig. 2.**
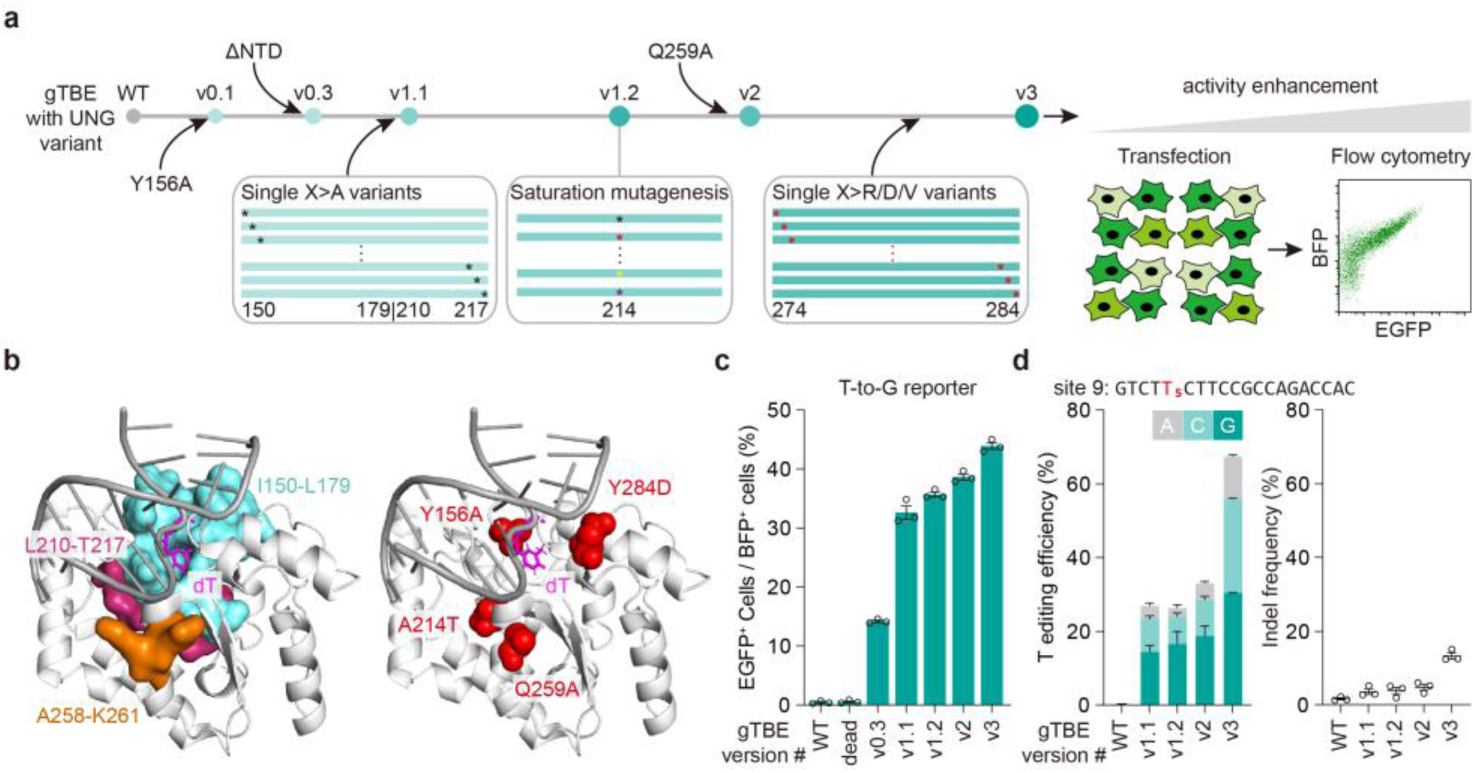
Protein engineering and evolution of gTBEs. **a**, Schematic diagram of mutagenesis and screening strategy for the engineered gTBE. The EGFP reporter plasmids were transiently co-transfected into cultured cells along with the gTBE plasmids, and the fluorescence intensity of EGFP was detected with flow cytometry. **b**, Left, the selected residues (shown as surface) for mutagenesis nearby the catalytic site pocket of human UNG-DNA complex (PDB entry 1EMH), in which dΨU was mutated to T in the DNA (dT). Right, location of the effective residues in gTBEv3 variant shown as spheres in red on the three-dimensional structure. **c**, Gradual improvement of EGFP activation for each gTBE variants (*n* = 3). WT, wild-type UNG2Δ88. dead, catalytically inactive UNG2Δ88 (carrying D154N and H277N mutations, equivalent to D145N and H268N of UNG1)^48^. **d**, Frequencies of T base editing outcomes (left) and indels (right) with different gTBE variants at the edited T5 position in site 9 in transfected HEK293T cells by target deep sequencing (*n* = 3). All values are presented as mean ±s.e.m.

Then, we examined the spatial neighbors of residue T214, nearby the Gly-Ser loop that compresses the DNA backbone 3′ to the lesion (Fig. 2b), and obtained variant gTBEv1.3 (v0.3 with Q259A), which increased the editing efficiency by 1.46-fold (Supplementary Fig. 4c). Furthermore, we found a synergistic enhancement of T-to-G editing activity in variant gTBEv2 (v0.3 with combination of A214T and Q259A), by 2.7-fold in comparison with the T editing activity of gTBEv0.3 (Fig. 2c). We also scanned residues in the regions of Q274-Y284, in or nearby the Leu-intercalation loop, by sequential replacement with amino acids of distinct properties, including arginine (with positive charged side chain), aspartic acid (with negative charged side chain), or valine (with small hydrophobic side chain) (X>R, D, or V). Although most of these mutations reduced the T editing activity, we found a variant gTBEv3 (v2 with Y284D) showed elevated editing efficiency by 1.22-fold as compared with that of gTBEv2 (Supplementary Fig. 5), and by 3.09-fold compared with gTBEv0.3 (Fig. 2c).

We validated the improvement of T editing activity by different gTBE variants at one endogenous genomic site in cultured mammalian cells (HEK293T). After transfected with all-in-one constructs encoding each gTBE variant, together with sgRNA that targeted site 9 and mCherry for fluorescence-activated cell sorting (FACS), mCherry-positive cells were FACS-sorted. Through target deep sequencing analysis, we obtained a gradual increase of overall T editing efficiency at T5 from 26.9% for gTBE1.1 to 67.4% for gTBE3, as well as the insertions and deletions (indels, from 3.6% to 13.3%), with T-to-S (i.e., T-to-C or T-to-G; S = C or G base) conversions as the predominant events at this site (Fig. 2d). These results indicate that rounds of mutagenesis described above had effectively optimized gTBE activity for T-to-C and T-to-G base editing. Thus, the engineered version of gTBEv3 (carrying Y156A, A214T, Q259A, Y284D mutations) had the highest T editing efficiency and was used for the following studies.

### Characterization of gTBEv3 at human genomic DNA sites

We further characterized the editing profiles of gTBEv3 by targeting 20 endogenous genomic loci, most of which were used in previous base editing studies^11, 12, 26, 27^. We found that gTBEv3 achieved efficient T base editing activity (ranged from 24.3% to 81.5%; Fig. 3a and Supplementary Fig. 6a,b), but essentially no A, C or G editing at all examined sites (Supplementary Fig. 6c-e). The T-to-C or T-to-G conversions were the predominant events (Supplementary Fig. 6f-h), with the ratios of T-to-S to T-to-A/C/G conversion ranged from 0.68 to 0.97 (Fig. 3b). Only a low percentage of T-to-A conversion were detected (Fig. 3a and Supplementary Fig. 6i), consistent with previous findings of gGBE^3^, AYBE^9^ and CGBEs^11–15^. We found that gTBEv3 also induced indels with frequency ranging from 5.2% to 45.2% at the 20 edited sites (Fig. 3c). Furthermore, we found that the editable range of gTBEv3 was positions 2 to 11, and the optimal editing window with high efficiency of T conversion covered protospacer positions 3 to 7, with the highest editing efficiency at position 5 (Supplementary Fig. 6b). We found no obvious motif preference for T conversions with gTBEv3 by analyzing the on-target editing and sequences of all tested sites (Supplementary Fig. 6j).

**Fig. 3.**
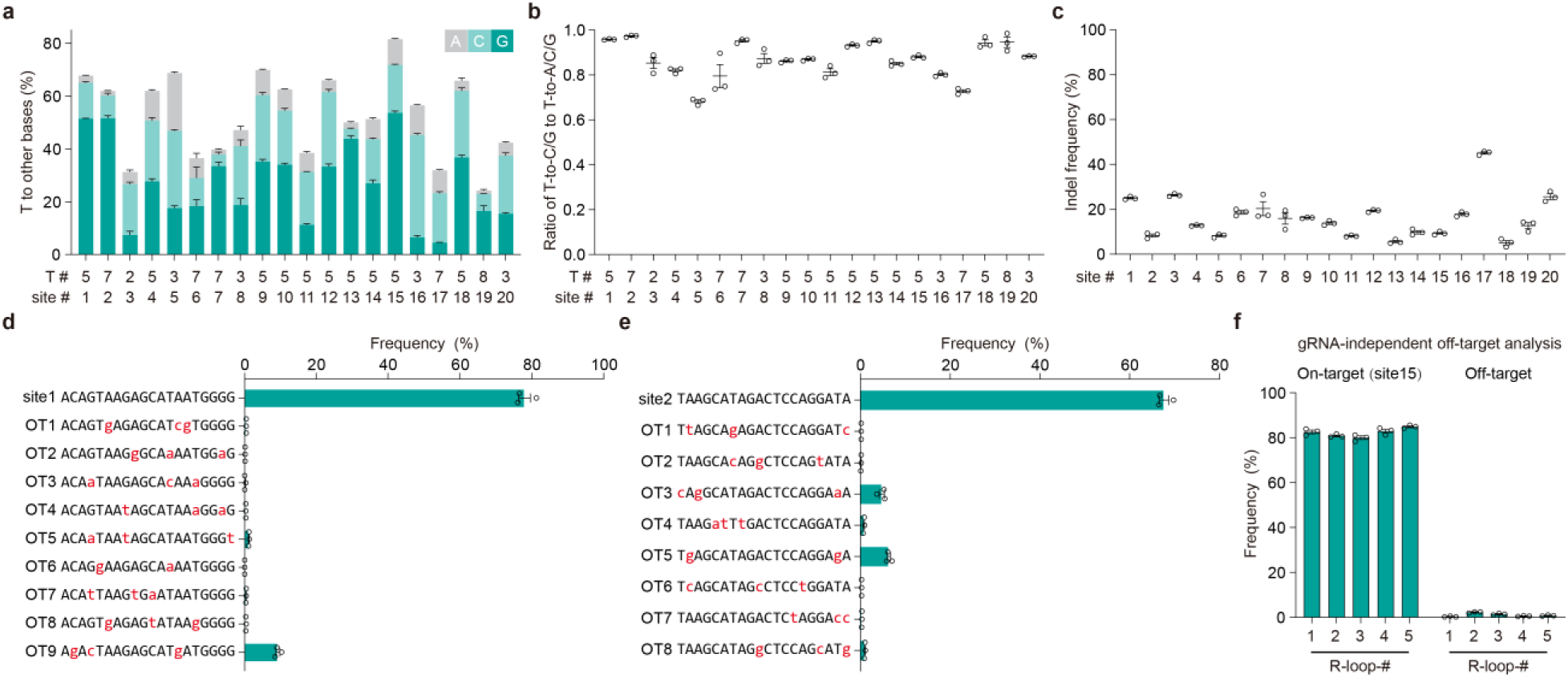
Characterization of editing profiles of gTBE via target deep sequencing. **a**, Bar plots showing the on-target DNA base editing at positions with the highest T conversion frequencies at each genomic site in HEK293T cells (mean ±s.e.m., *n* = 3). T#: T position with highest on-target base editing frequencies across protospacer positions 1–20. site #: genomic site number. **b**, The ratio of T-to-C/G to T-to-A/C/G conversion frequency by gTBEv3 editing at the sites shown in **a**. **c**, indels frequencies with gTBEv3 at 20 on-target sites (*n* = 3). **d**,**e**, The sgRNA-dependent off-target analysis for gTBEv3 editing efficiency at site 1 and site 15 (*n* = 3). OT: off-target. **f**, The sgRNA-independent off-target editing efficiency detected by the orthogonal R-loop assay at each R-loop site (*n* = 3). All values are presented as mean ±s.e.m.

We have analyzed the off-target activity of gTBEv3 at several in silico-predicted^28^ guide-dependent off-target sites, and characterized the ability of gTBEv3 to mediate guide-independent off-target DNA editing using orthogonal R-loop assay in five previously reported dSaCas9 R-loops^9, 29^. We found very low percentage of editing at all the guide-dependent off-target loci (Fig. 3d,e and Supplementary Fig. 7) and detected very low frequencies (1.1% in average) at all five guide-independent off-target sites (Fig. 3f). Taken together, the gTBEv3 represents a highly efficient T-to-S base editor with low off-target effects in mammalian cells.

### Enhancement of C editing activity of gCBE

To examine whether the mutations emerged from the engineering of gTBE would benefit the enhancement of gCBE activity, we attempted to introduce A214V into gCBEv0.3 using the C-to-G reporter to evaluate the editing activity (Fig. 4a). The resulted gCBEv1.1 largely elevated the C-to-G conversion activity by 1.34-fold (Supplementary Fig. 8a). We conducted alanine-scanning mutagenesis on the fragment of D154-D189 to examine its role in the regulation of base excision activity, and obtained a variant gCBEv1.2 (v0.3 with K184A) largely elevating the C-to-G conversion activity by 1.55-fold (Supplementary Fig. 8b). We further investigated the additive effect of A214V and K184A by combining these two mutations in gCBEv2 (carrying K184A, N213D, A214V mutations), and found synergistic enhancement of C-to-G editing activity by 1.3-fold compared with that of gCBEv0.3 (Fig. 4b). We further validated the improvement of C editing activity for different gCBE variants by targeting an endogenous genomic site, and found a gradual increase of overall C editing efficiency from 18.2% to 37.2% at C2 of the site 28 (Supplementary Fig. 9a).

**Fig. 4.**
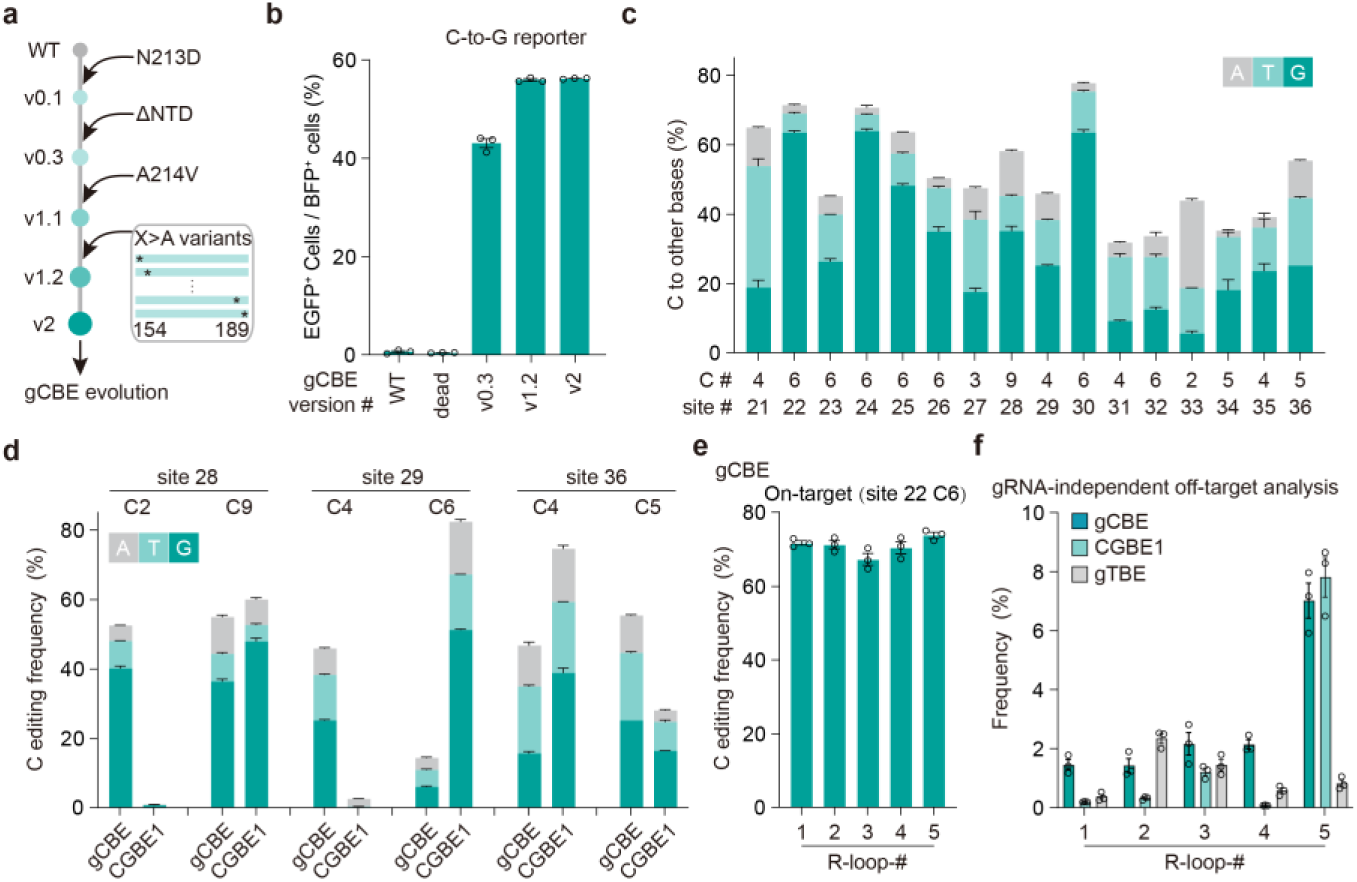
Enhancement of gCBE editing activity through protein engineering. **a**, Schematic diagram of mutagenesis and screening strategy for the engineered gCBE. **b**, Gradual improvement of EGFP activation for each gCBE variants (*n* = 3). WT, wild-type UNG2Δ88. dead, catalytically inactive UNG2Δ88 (carrying D154N and H277N mutations, equivalent to D145N and H268N of UNG1)^48^. **c**, Bar plots showing the on-target DNA base editing at positions with the highest C conversion frequencies at each genomic site in HEK293T cells (*n* = 3). C#: C position with highest on-target base editing frequencies across protospacer positions 1-20. site #: genomic site number. **d**, Bar plots showing the on-target DNA base editing of different positions at three loci with gCBE or CGBE1 (*n* = 3). **e**, On-target base editing frequencies for gCBEv2 at C6 of site 22 in HEK293T cells for the orthogonal R-loop assay (*n* = 3). **f**, gRNA-independent off-target editing frequencies detected by the orthogonal R-loop assay at each R-loop site. Each R-loop was performed by co-transfection of each base editor, and an SpCas9 sgRNA targeting corresponding site with dSaCas9 and a SaCas9 sgRNA (*n* = 3). All values are presented as mean ±s.e.m.

By targeting 16 endogenous genomic loci, we characterized the editing profiles of gCBEv2 and obtained efficient C base editing activity ranged from 31.8% to 77.7% (Fig. 4c and Supplementary Fig. 9b-d). We found that gCBEv2 could induce predominant C-to-G conversions as well as C-to-T conversions, with the ratios of C-to-G/T to C-to-A/G/T conversion reaching up to 0.97, and there were very few C-to-A conversions detected (Fig. 4c, Supplementary Fig.9e-h). The gCBEv2 could induce indels with frequency ranged from 3.1% to 48.3% at the examined sites (Supplementary Fig.9i). After analyzing the sequences of all tested sites, we found that the editable range of gCBEv2 was positions 2 to 9 (Supplementary Fig. 9c), and gCBEv2 showed preferences for editing at AC or TC motifs with a higher efficiency than other motifs (Supplementary Fig. 9j).

When compared to CGBE1^12^, a C-to-G base editor, we found that gCBEv2 showed higher editing activity at certain positions towards the distal end of the target sequence (Fig. 4d and Supplementary Fig. 9c), indicating their positional preferences within different optimal editing windows (positions 2 to 6 for gCBEv2 v.s. positions 5 to 7 for CGBE1^12^). The gCBEv2 induced fewer indels at site 36, and more indels at site 28 and site 29 than CGBE1 (Supplementary Fig.9k). Using the orthogonal R-loop assay^9, 29^ mentioned above, we characterized comparably low frequencies (< 2% in average) at 4/5 guide-independent off-target sites (Fig. 4e,f and Supplementary Fig. 9l).

Moreover, we found that the gCBEv2 could only facilitate C editing, but there was essentially no T editing at all examined sites (Supplementary Fig. 9c,d). The editing specificity of gCBEv2, together with that of gTBEv3 (Supplementary Fig. 6b-e), consolidated the orthogonality of these two base editors for base editing.

### Applications of gTBE and gCBE

We further evaluated the potential applications of gTBE and gCBE. The gTBE could not only remediate inactive splicing signals in the intron-split EGFP reporter systems used above (Fig. 1, 2 and Supplementary Fig. 2), but also be used for exon skipping by disrupting splicing signals at splicing donor (SD) or splicing acceptor (SA) sites (Fig. 5a). After analyzing the splicing sites in 16 well-studied genes for gene and cell therapy research^30–32^, we found that gTBE and gCBE, together with other existing base editors, provide 1904 sgRNA candidates (Supplementary Table 4) with the SD or SA sites located in each optimal editing window (Fig. 5b and Supplementary Fig. 10a). Among the 771 sgRNA candidates for ABE and CBE targeting, 156 and 103 candidates overlapped with those for gGBE and gTBE, respectively (Fig. 5c). Moreover, 232 and 223 sgRNA candidates could only be screened by gGBE or gTBE targeting, respectively (Fig. 5c). For gCBE, apart from 205 sgRNA candidates overlapped with those for CBE, there were 148 unique candidates (Supplementary Fig. 10b). The availability of these base editors could largely expand the scope of sgRNA screening for efficient editing at splicing sites (Supplementary Fig. 10). In addition, the newly developed base editors could be utilized for bypassing premature termination codons (PTCs) and introduction of PTCs (Supplementary Fig. 11). The gTBE and gCBE could provide more versatile codon outcomes from PTCs editing (Supplementary Fig. 11b), and introduce PTCs by editing more codons coding various amino acids (Supplementary Fig. 11d). To potentially disrupt gene function by introduction of PTCs, we analyzed and obtained 851 sgRNA candidates (Supplementary Table 5) targeting various codons for PTCs introduction in 15 genes with gGBE and CBE, with 191 TAC and 124 TCA for gGBE targeting (Supplementary Fig. 11e).

**Fig. 5.**
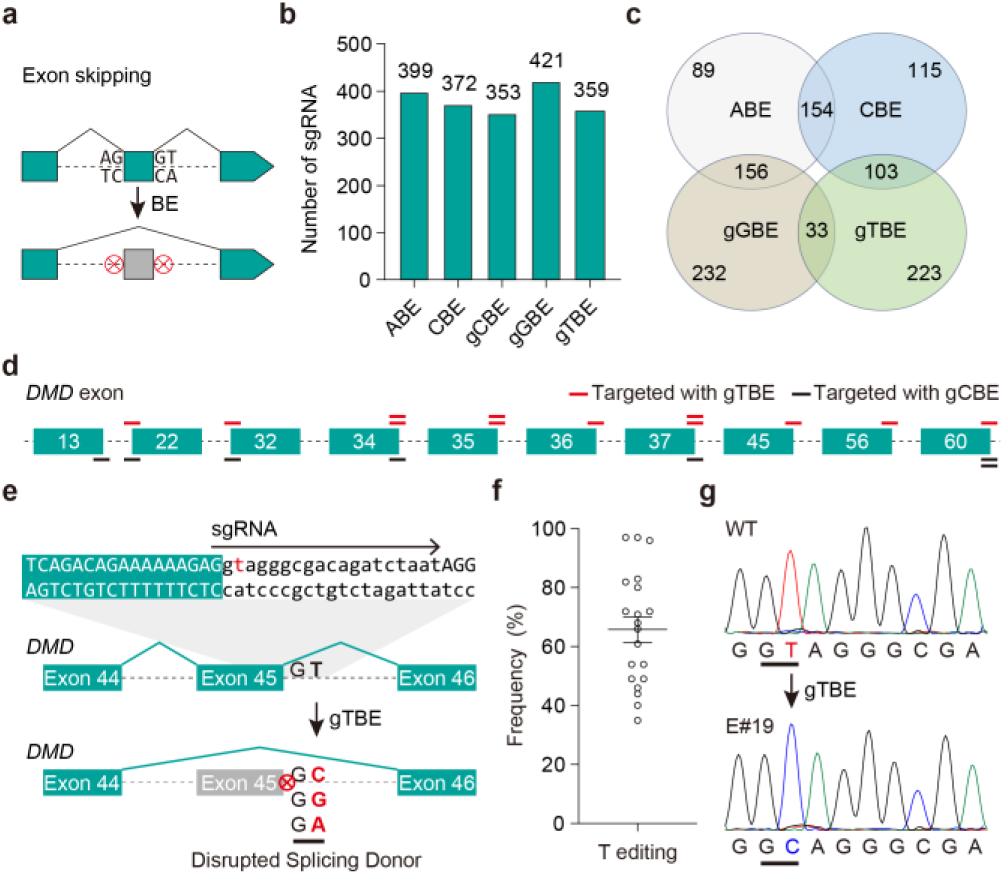
Gene editing applications of gTBE and gCBE. **a**, principle for exon skipping with base editors. **b**, Bar plots showing the numbers of sgRNA candidates targeting the splicing sites in 16 genes by different base editors. The 16 genes are *AGT, ANGPTL3, APOC3, B2M, CD33, DMD, DNMT3A, HPD, KLKB1, PCSK9, PDCD1, PRDM1, TGFBR2, TRAC, TTR,* and *VEGFA*. **c**, Venn diagram showing the distribution of sgRNAs for 4 base editors in **b**. **d**, Schematic diagram illustrating sgRNA candidates specifically targeting SD or SA sites in human *DMD* with gTBEv3 (red lines) or gCBEv2 (black lines), but not ABE or CBE. **e**, Schematic diagram illustrating the skipping of human *DMD* exon 45 induced by gTBE-induced disruption of the splicing donor site. **f**, On-target base editing efficiency for gTBEv3 targeting the splicing donor site of humanized *DMD* exon 45 in mouse embryos (mean ±s.e.m., *n* = 20). Sanger sequencing results were quantified by EditR. **g**, DNA sequencing chromatograms from wild-type (WT) and representative embryos co-injected with gTBEv3 mRNA and sgRNA targeting the SD site of human *DMD* exon 45.

To illustrate these applications, we focused on editing the splicing sites in human *DMD* gene (*Duchenne muscular dystrophy*, coding dystrophin) that could not be targeted with ABE or CBE. We designed and screened a series of sgRNAs specifically targeting SD or SA sites with gTBEv3 or gCBEv2 (Fig. 5d and Supplementary Fig. 10c), including two sgRNAs targeting the SD site of *DMD* exon 37 (Supplementary Fig. 10d) and exon 45 (Fig. 5e) uniquely by gTBEv3. Disruption of the SD site of exon 45, thus leading to exon skipping, would be applicable to restore dystrophin expression in 9% DMD patients^33^. After co-injection of gTBEv3 mRNA and sgRNA targeting the SD site of *DMD* exon 45 into zygotes of humanized mice, 100% (20/20) mouse embryos harbored efficient base conversion (ranged from 35.0% to 97.0%) at the desired position T3 (Fig. 5f,g), indicating the great potential of gTBE for human disease modeling and gene therapy. Overall, gBEs, including gTBE, gCBE and gGBE, provide more options for the sites that dBEs could not target, largely expanding the targeting scope of base editors.

## Discussion

The deaminase-based base editor (dBE) and deaminase-free glycosylase-based base editor (gBE) are currently two main categories of DNA base editors^3^, enabling direct editing of adenine (A), cytosine (C), or guanine (G), but not thymine (T). In this study, we engineered two orthogonal base editors, gTBE and gCBE, that could achieve highly efficient T and C editing in both cultured human cells and mouse embryos. We have shown that the same original DNA glycosylase could be engineered into enzymes that selectively excise specific nucleotide bases and harnessed to develop novel base editors using the deaminase-free glycosylase-based strategy. The enhanced editing efficiency could be attributed to mutations in the UNG moiety that facilitate its specific substrate preference or ssDNA-binding activity, or both, which needs to be elucidated by biochemical and structural experiments in the future. Although our mutagenesis and screening strategy based on rational design was effective, the mutagenesis was far from saturating the potential mutant repertoire. More other mutations in other positions of UNG would be identified to enhance the editing activity of gTBE and gCBE.

In human, about 19% of the pathogenic single nucleotide polymorphism (SNP) could be corrected through T-to-G conversion^9^. The gTBE and gCBE could greatly broaden the targeting scope of base editors by breaking the limitations of PAM and narrow editing window, thus increasing the opportunity to obtain an efficient strategy for further research. The T-to-S conversion ability of gTBE allows for a variety of gene-editing applications, including editing splicing sites, as well as editing that bypass PTCs. Although we have evaluated the off-target effects of gTBE and gCBE on several targeted genes, a comprehensive analysis through high-throughput whole-genome sequencing methods, such as GOTI^34^ and SAFETI^35^, is required for a thorough assessment of off-target effects before their potential therapeutic applications. Although the editable range (positions 2 to 11) is wide with the current version of gTBE (Supplementary Fig. 6b), a more accurate gTBE with a refined editing window might be achieved through further engineering of the glycosylase moiety or architectures of gTBE, encouraged by the development of ABE9^36^ or YE1-BE3^37^. Moreover, more specific T-to-C, T-to-G, or C-to G editors could potentially be achieved by harnessing the DNA repair machinery in the base excision repair (BER) pathway^9, 38–42^ or by further structural fine-tuning of gTBE or gCBE. Due to one-step generation of AP sites by engineered glycosylases, gBEs facilitating A or C editing might have great potential advantages over certain dBEs (e.g., AYBE, CGBE). We note that indels induced by gTBE and gCBE, as well as by AYBE, AXBE and CGBEs that generating AP sites, were higher than those by ABE or CBE^4, 5, 9–15^. Encouraged by the previous studies on CGBE^12, 15^ and AYBE^9^, additional effort is required to further reduce the level of off-target editing or indels through engineering approaches.

In summary, we have engineered two orthogonal base editors based on the same original DNA glycosylase for direct T editing and C editing. These findings also indicated that other DNA glycosylases could be engineered into enzymes that selectively excise specific nucleotide bases and be applied to develop novel base editors with various features, thus broadening the editing scope of base editors.

To be noted, when we were preparing this manuscript, Chang and colleagues preprinted their work reporting base editor TSBE for T-to-G base editing developed using protein language models^43^. The results convinced that direct T editing can be achieved using the deaminase-free glycosylase-based strategy by engineering certain uracil DNA glycosylase. Although the performance of TSBE was improved, gTBE showed higher editing efficiency than TSBE (Supplementary Fig. 12). In contrast to protein language models, we rationally designed and screened 91 and 37 UNG variants for engineering gTBEv3 and gCBEv2, respectively, and obtained marked enhancement of base editing activity. In addition, the TSBEs were developed using the full length UNG2, in which the disordered N-terminal domain (NTD) containing motifs and sites for undesired protein-protein interactions and post-translational modifications^19, 20^. Since the N-terminus plays important roles in enhancing the translocation of hUNG2 on DNA^18^, TSBEs might induce more undesired edits and await further characterization before widespread use.

## Acknowledgments

This work was supported by HuidaGene Therapeutics Co., Ltd. (H.T.).

## Author contributions

H.Y. and H.T. jointly conceived the project. H.T. designed and conducted experiments. H.W. and N.L. performed experiments with the help of Y.L., D.W. and X.W.; G.L., M.J., H.L. and Y.W. performed mouse experiments; Y.Z. performed bioinformatics analysis. Y.Y. assisted with data analysis. H.Y., L.S., X.Y. and H.T. supervised the whole project. H.Y. and H.T. wrote the manuscript, and all authors contributed to the editing of the manuscript.

## Methods

### Molecular cloning

Base editor constructs used in this study were cloned into a mammalian expression plasmid backbone under the control of a EF1α promoter by standard molecular cloning techniques, and the two intron-split EGFP reporters were constructed similar to those described previously^9^, except that the engineered sequence containing the last 86 base pairs (bp) intron of human *RPS5* was inserted between BFP and EGFP coding sequences. And the corresponding mutations at the splice acceptor site were made to construct T-to-G reporter or C-to-G reporter via site-directed mutagenesis by PCR, respectively. Mutations at the splice acceptor site led to inactive EGFP production. Encouraged by the findings from previous base editors^12, 15^, the corresponding mutations at the splice acceptor site were put at position 6 across the protospacer.

The wild-type UNG2 sequence (313 amino acids long) was PCR-amplified from cDNA of HEK293T, and fused at the C-terminus of nCas9 via Gibson Assembly method. UNG2-Y156A, UNG2-N204D, BpiI-harboring mutants, UNG-NTD-truncated mutants and corresponding combinations were constructed via site-directed mutagenesis by PCR. UNG mutagenesis libraries were designed and generated as previously described^44^. The selected regions of UNG1 were divided into 8 aa long segments. Sequential alanine / arginine / aspartic acid / valine substitutions (X>A, R, D, or V) and site-saturation mutagenesis of the residue 214 were designed, with oligos coding for the mutants annealed and ligated into corresponding BpiI-digested backbone vectors.

The Cas-OFFinder^28^ was used to search for potential guide-dependent off-target sites of Cas9 RNA-guided endonucleases with a maximum of 3 mismatches (with no bulges). For sgRNAs targeting *DMD* splicing sites with an NGN PAM, a PAM-flexible Cas9 variant SpG was used. The sgRNA oligos were annealed and ligated into BpiI sites. The amino-acid sequence for gTBEv3 was supplied in Supplementary Table 1. The UNG mutants and corresponding codon substitutions used were listed in Supplementary Table 2.

### Cell culture, Transfection, and flow cytometry analysis

HEK293T cells were cultured with DMEM (Catalog# 11995065, Gibco) supplemented with 10% fetal bovine serum (Catalog# 04-001-1ACS, BI) and 0.1 mM non-essential amino acids (Catalog# 11140-050, Gibco) in an incubator at 37 °C with 5% CO_2_.

Mutant screening was conducted in 48-well plates, with 3 ×10^4^ HEK293T cells per well plated in 250 μL of complete growth medium the day before transfection. Between 16 and 24 h after seeding, cells were co-transfected with 250 ng gTBE (or gCBE) plasmids, 250 ng T-to-G (or C-to-G) reporter plasmids and 1 μg Polyethylenimine (PEI) (DNA/PEI ratio of 1:2) per well. For cell transfection of HEK293T for FACS, 5 ×10^5^ cells per well were plated in 12-well plates with 1 ml complete growth medium the day before transfection. After 14-16h, 2 μg all-in-one plasmids containing gTBE and corresponding sgRNA were transfected into cells using PEI (DNA/PEI ratio of 1:2). Orthogonal R-loop assays were performed as described previously^9, 29^. In brief, 1 μg of gTBE or gCBE plasmid with sgRNA targeting the corresponding site (with mCherry as reporter) and 1 μg of dSaCas9 plasmid with corresponding sgRNA targeting five off-target sites to generate R-loops (with EGFP as reporter) were co-transfected into HEK293T cells in 12-well plates using PEI (DNA/PEI ratio of 1:2).

At 48h post-transfection, expression of mCherry, BFP and EGFP fluorescence were analyzed by BD FACS Aria III or Beckman CytoFLEX S. Flow cytometry results were analyzed with FlowJo V10.5.3. The gating strategy in the identification of mCherry^+^, BFP^+^ and EGFP^+^ cells for on-target editing efficiency evaluation was supplied in Supplementary Figure 2b.

### Target sequencing of endogenous sites and data analysis

Endogenous target sites of interest were amplified from genomic DNA as previously described^9^. Briefly, 10,000 positive cells with mCherry were isolated by FACS after 72 h of transfection, then genomic DNA was extracted and the regions of interest for target sites were amplified by PCR using site-specific primers. The purified PCR products were analyzed by Sanger sequencing (Genewiz).

Target sequencing data analysis was described in the previous paper^3^. In brief, the amplicons were ligated to adapters and sequencing was performed on the Illumina MiSeq platforms, then the targeted amplicon sequencing reads were processed using fastp with default parameters^45^, and further amplicon sequencing analysis were performed by CRISPResso2^46^. T-to-G purity was calculated as T-to-G editing efficiency / (T-to-C editing efficiency + T-to-G editing efficiency + T-to-A editing efficiency). T-to-S conversion ratio was calculated as (T-to-C editing efficiency + T-to-G editing efficiency) / (T-to-C editing efficiency + T-to-G editing efficiency + T-to-A editing efficiency). In Figure 5e, the frequencies of editing were calculated from Sanger sequencing traces using EditR^47^ (https://moriaritylab.shinyapps.io/editr_v10/). Protospacer sequences and site-specific primers used for each genomic locus are listed in Supplementary Table 3.

### In *vitro* transcription of gTBE mRNA and *DMD* sgRNA

The mRNA and sgRNA preparations were performed as previously described^9^. The gTBE plasmids were linearized by the FastDigest KpnI restriction enzyme (Catalog# FD0524, Thermo Fisher), purified using Gel Extraction Kit (Catalog# D2500-03, Omega), and used as the template for *in vitro* transcription (IVT) using the mMESSAGE mMACHINE T7 Ultra kit (Catalog# AM1345, Thermo Ambion). For *DMD*-sgRNA preparation, we added the T7 promoter sequence to the sgRNA template by PCR amplification. The T7-*DMD*-sgRNA PCR product was purified using Gel Extraction Kit (Catalog# D2500-03, Omega) and used as the template for IVT of sgRNAs using the MEGAshortscript T7 kit (Catalog# AM1354, invitrogen). The gTBE mRNA and *DMD*-sgRNA were purified using the MEGAclear kit (Catalog# AM1908, invitrogen), eluted in RNase-free water and stored at -80°C until use.

### Animals and microinjection of mouse zygotes

Animal manipulations were consistent with those reported previously^3^. Experiments involving mice were approved by the Biomedical Research Ethics Committee of Center for HuidaGene Therapeutics Co. Ltd. Mice were maintained in a specific pathogen-free facility under a 12-hour dark–light cycle, and constant temperature (20–26°C) and humidity (40–60%) maintenance.

Super ovulated humanized *DMD* females with human *DMD* exon 45 in C57BL/6 background (4 weeks old) were mated with C57BL/6 males (8 weeks old), and females from the ICR strain were used as foster mothers. Fertilized embryos were collected from oviducts 21 h post hCG injection. For zygote injection, the mixture of gTBE mRNA (250 ng/µL) and *DMD*-sgRNA (250 ng/µL) was injected into the cytoplasm of 1-cell embryo in a droplet of M2 medium using a FemtoJet microinjector (Eppendorf) with constant flow settings. The injected embryos were cultured in M16 medium with amino acids to blastocysts for three days (37°C and 5% CO2) before genomic DNA extraction and target amplification.

### Statistical analysis

Statistical tests performed by Graphpad Prism 8 included the one-tailed paired two-sample *t*-test or Dunnett’s multiple comparisons test after one-way ANOVA.

## Data availability

All data supporting the findings of this study are available in the paper (and in its supplementary information files). Targeted amplicon sequencing data have been deposited at the Sequence Read Archive. All relevant original data are available from the corresponding authors upon request.

## Code availability

Custom scripts for CRISPResso analyses supporting the findings of this study are available from the corresponding author upon reasonable request.

## Supplementary figures

**Supplementary Figure 1.**
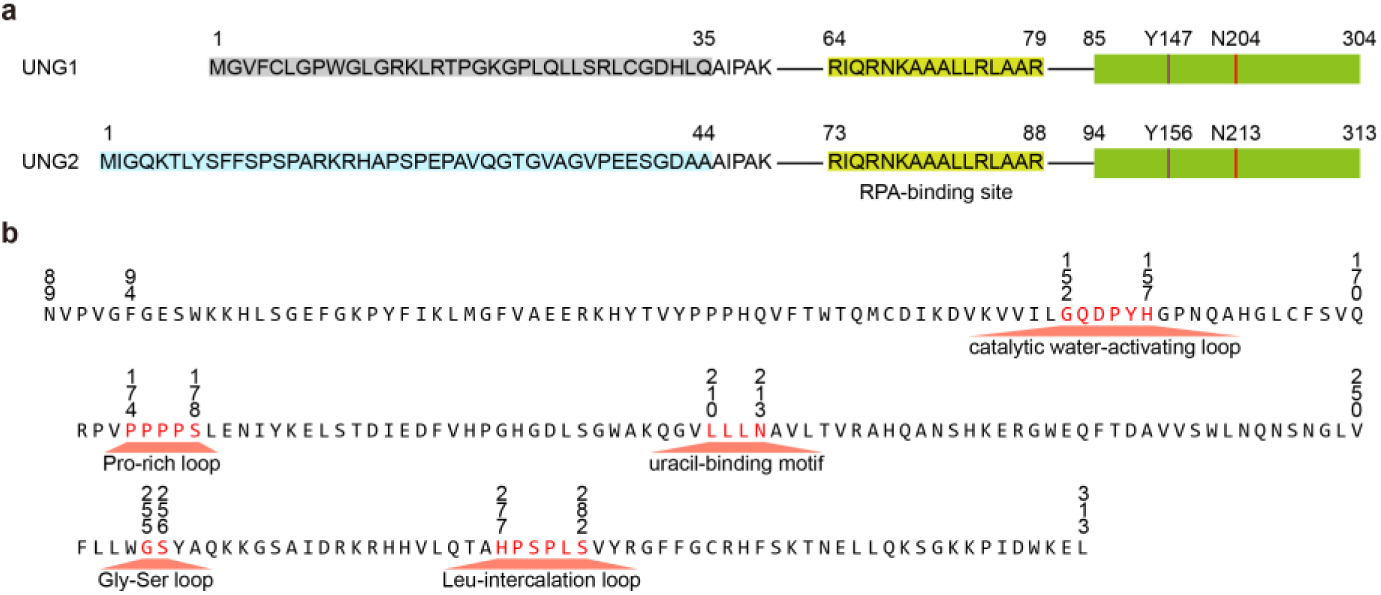
Characteristic sequences and motifs of human UNG1 and UNG2. **a**, UNG1-specific N-terminal residues (amino acid 1-35) are marked in grey. UNG2-specific N-terminal residues (amino acid 1-44) are light blue. The common RPA-binding site (yellow) and the globular catalytic domain (light green) are indicated. RPA, Replication protein A. **b**, UNGs contain five conserved motifs numbered from UNG2 as follows: the catalytic water-activating loop (152-GQDPYH-157); the proline (Pro) -rich loop compressing the DNA backbone 5’ to the lesion (174-PPPPS-178); the uracil-binding motif (210-LLLN-213); the glycine-serine (Gly-Ser) loop that compresses the DNA backbone 3’ to the lesion (255-GS-256); and the leucine (Leu) - intercalation loop penetrating the minor groove (277-HPSPLS-282).

**Supplementary Figure 2.**
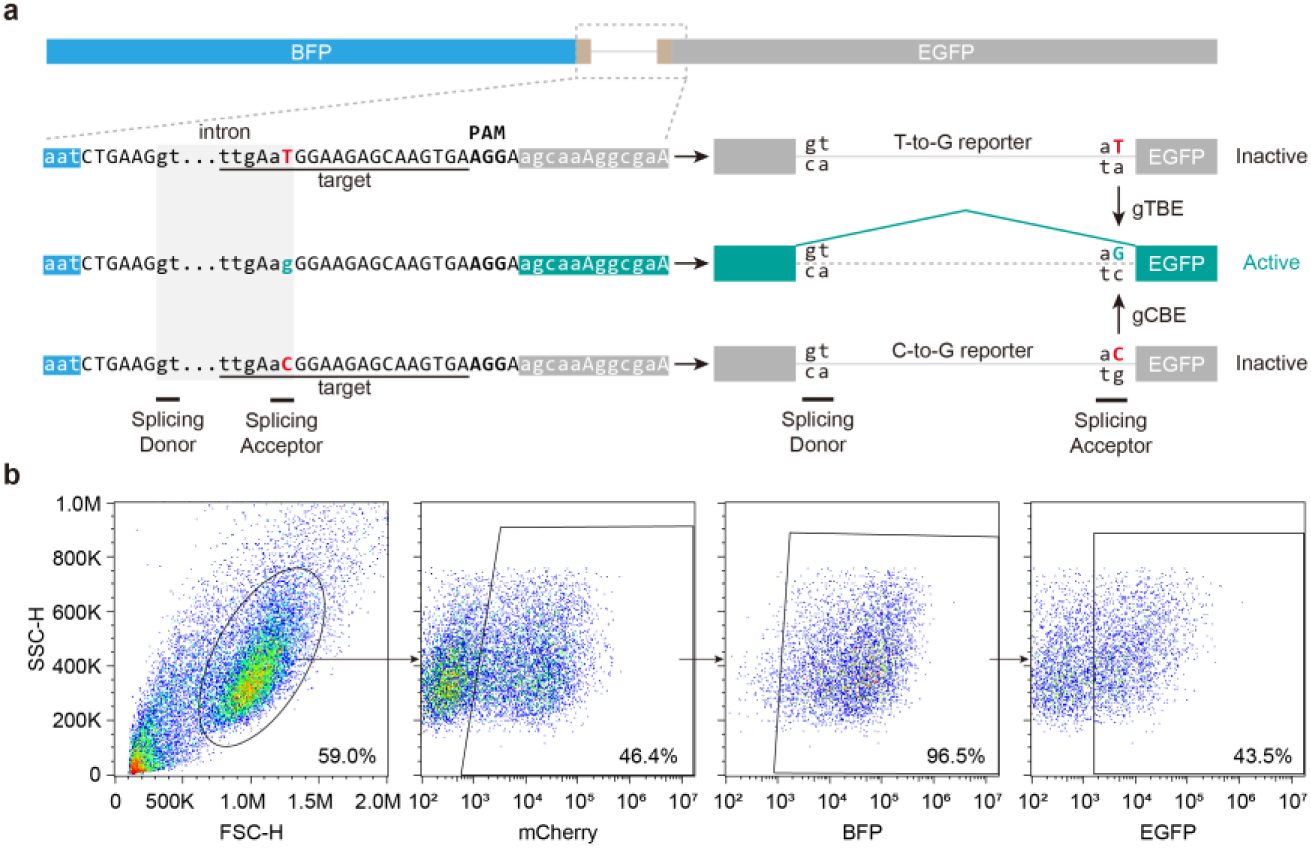
Characterizations of T-to-G and C-to-G reporter system. **a**, Schematic construct designs of the reporter for T-to-G or C-to-G editing detection. PAM, Protospacer adjacent motif. **b**, Representative flow cytometry scatter plots showing gating strategy and the percentages of gated cells for gCBEv0.3.

**Supplementary Figure 3.**
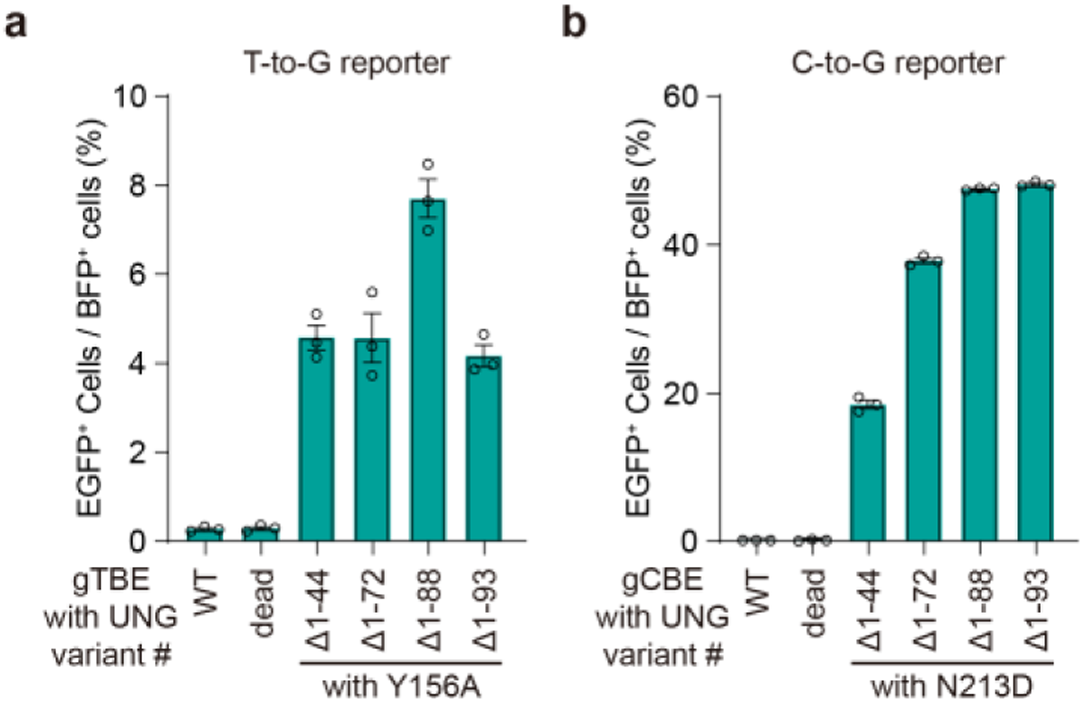
Editing activity of gTBE and gCBE candidates with various UNG-NTD truncations. **a**, Percentage of EGFP^+^ cells for evaluating T editing activity of gTBE candidates with various UNG variants (*n* = 3). **b**, Percentage of EGFP^+^ cells for evaluating C editing activity of gCBE candidates with various UNG variants (*n* = 3). WT, wild-type UNG2Δ88. dead, catalytically inactive UNG2Δ88 (carrying D154N and H277N mutations, equivalent to D145N and H268N of UNG1). All values are presented as mean ±s.e.m.

**Supplementary Figure 4.**
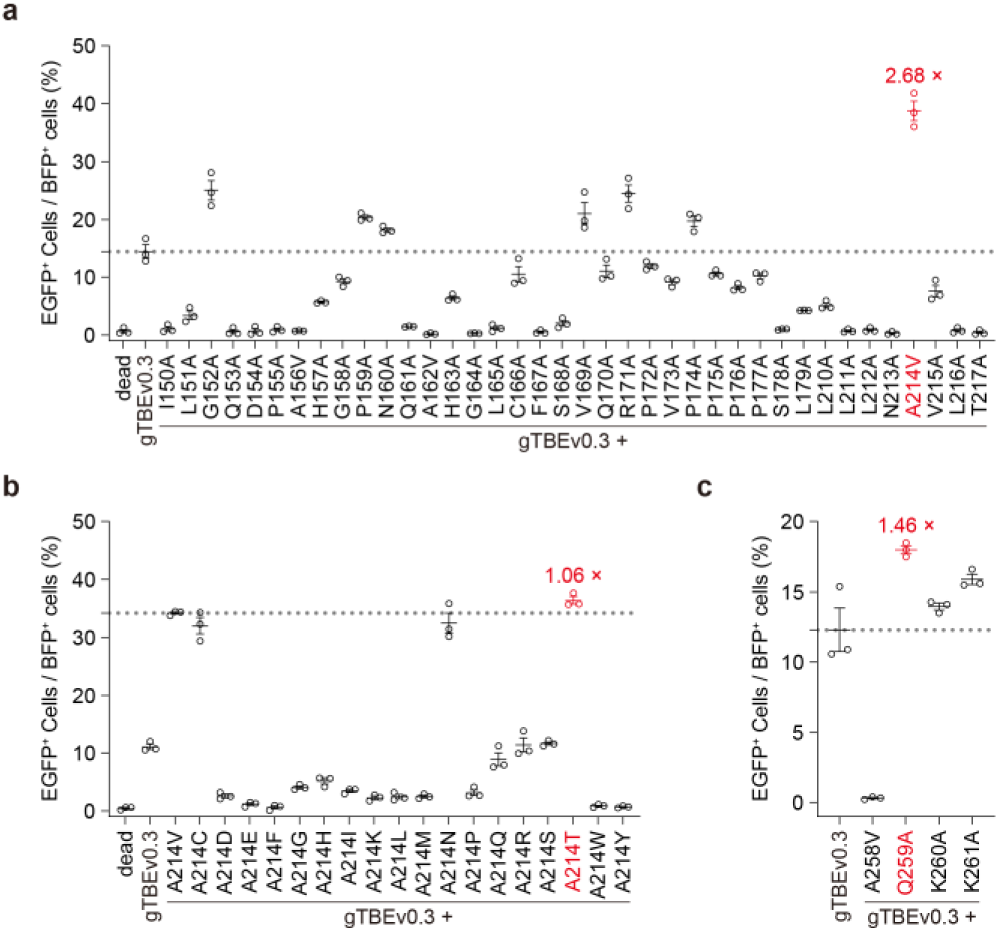
Performance of engineered variants in the gTBEv0.3 background. **a**, Percentage of EGFP^+^ cells of gTBE variants from alanine-scanning mutagenesis of regions covering the catalytic water-activating loop, the Pro-rich loop, and the uracil-binding motif (*n* = 3). Replacement of alanine with valine, A>V, is intended to cover all the residues in the interested regions. **b**, Percentage of EGFP^+^ cells of gTBE variants from site-saturation mutagenesis of the residue at position 214 (*n* = 3). **c**, Percentage of EGFP^+^ cells of gTBE variants with mutations of selected spatial neighbors of residue T214 (*n* = 3). dead, catalytically inactive UNG2Δ88 (carrying D154N and H277N mutations, equivalent to D145N and H268N of UNG1). All values are presented as mean ±s.e.m.

**Supplementary Figure 5.**
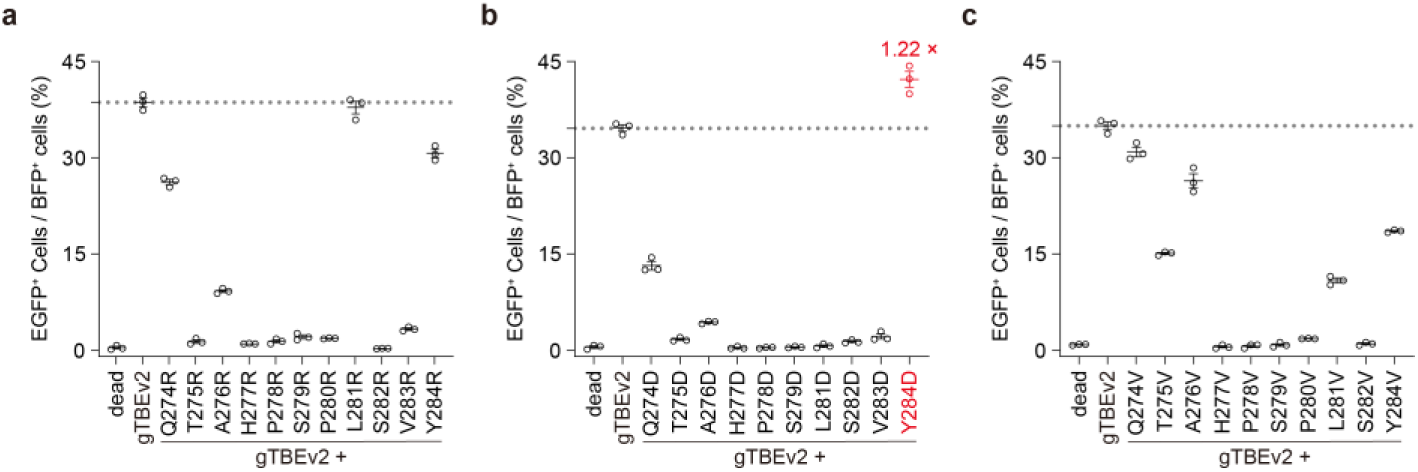
Performance of engineered variants in the gTBEv2 background. **a-c**, Percentage of EGFP^+^ cells of gTBE candidates with different UNG variants from sequential substitutions of arginine (**a**), aspartic acid (**b**), and valine (**c**) (X>R, D, or V) (*n* = 3). dead, catalytically inactive UNG2Δ88 (carrying D154N and H277N mutations, equivalent to D145N and H268N of UNG1). All values are presented as mean ±s.e.m.

**Supplementary Figure 6.**
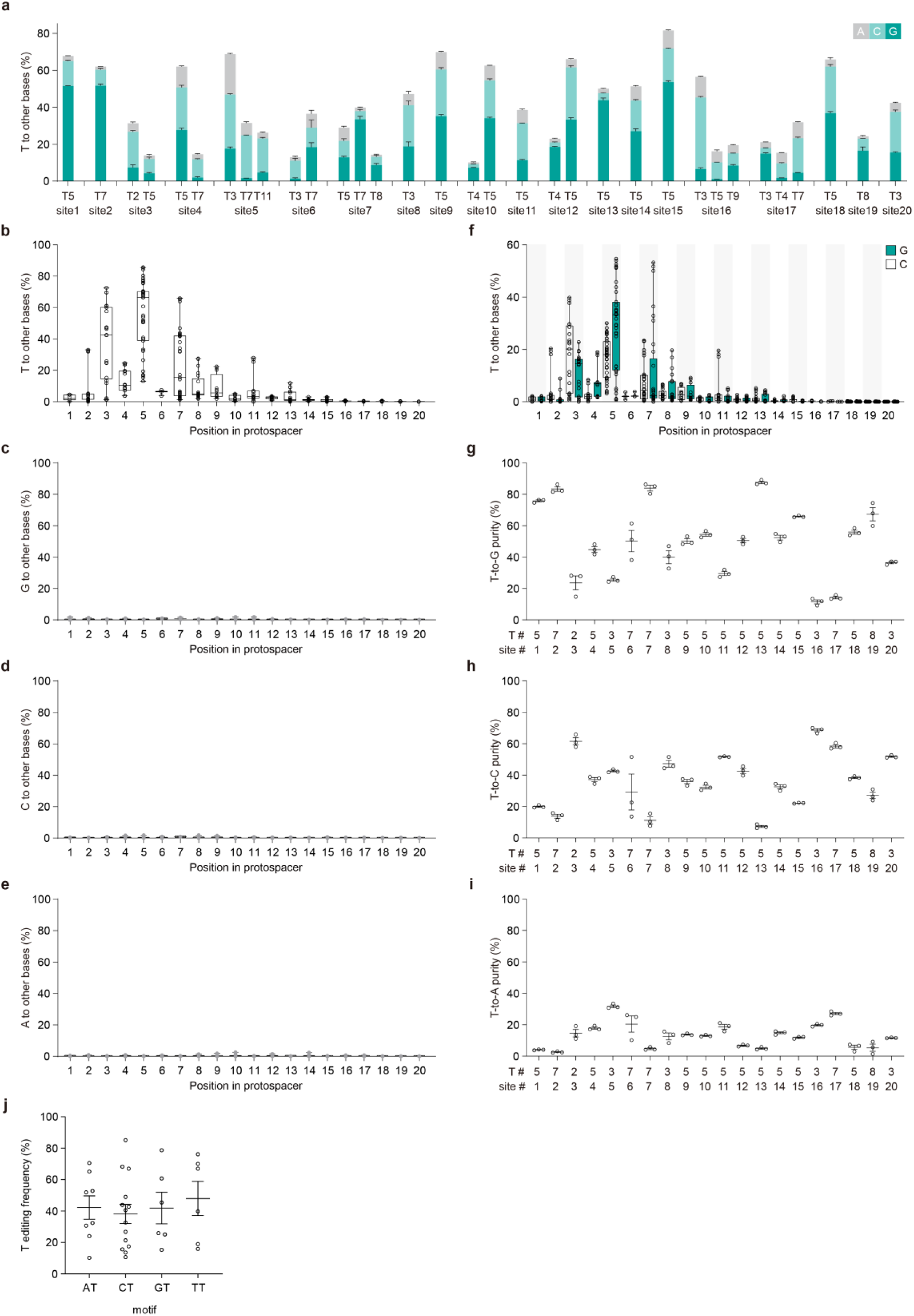
Further characterization of editing profiles for gTBEv3. **a**, Bar plots showing the on-target DNA base editing at positions with T conversion frequencies >10% at each genomic site in HEK293T cells (mean ±s.e.m., *n* = 3). **b-e**, Frequencies of T (**b**), G (**c**), C (**d**) and A (**e**) conversions by gTBEv3 across the protospacer positions 1-20 (where PAM is at positions 21–23) from the edited sites in Figure 3a. **f**, Frequencies of T-to-G and T-to-C editing by gTBEv3. In **b-f**, single dot represents individual replicate (*n* = 3 independent replicates per site), and boxes span the interquartile range (25th to 75th percentile); horizontal lines within the boxes indicate the median (50%); and whiskers extend to the minimal and maximal values. **g-i**, Percentage of T-to-G (**g**), T-to-C (**h**) or T-to-A (**i**) editing by gTBEv3 at various edited sites shown in Figure 3a (mean ±s.e.m., *n* = 3). T#: T position with highest on-target base editing frequencies across protospacer positions 1–20. site #: genomic site number. **j**, The statistical analysis of on-target DNA base editing for each NT motif from the edited sites in (**a**). Each dot represents the mean of three biological replicates for each edited position at various edited sites.

**Supplementary Figure 7.**
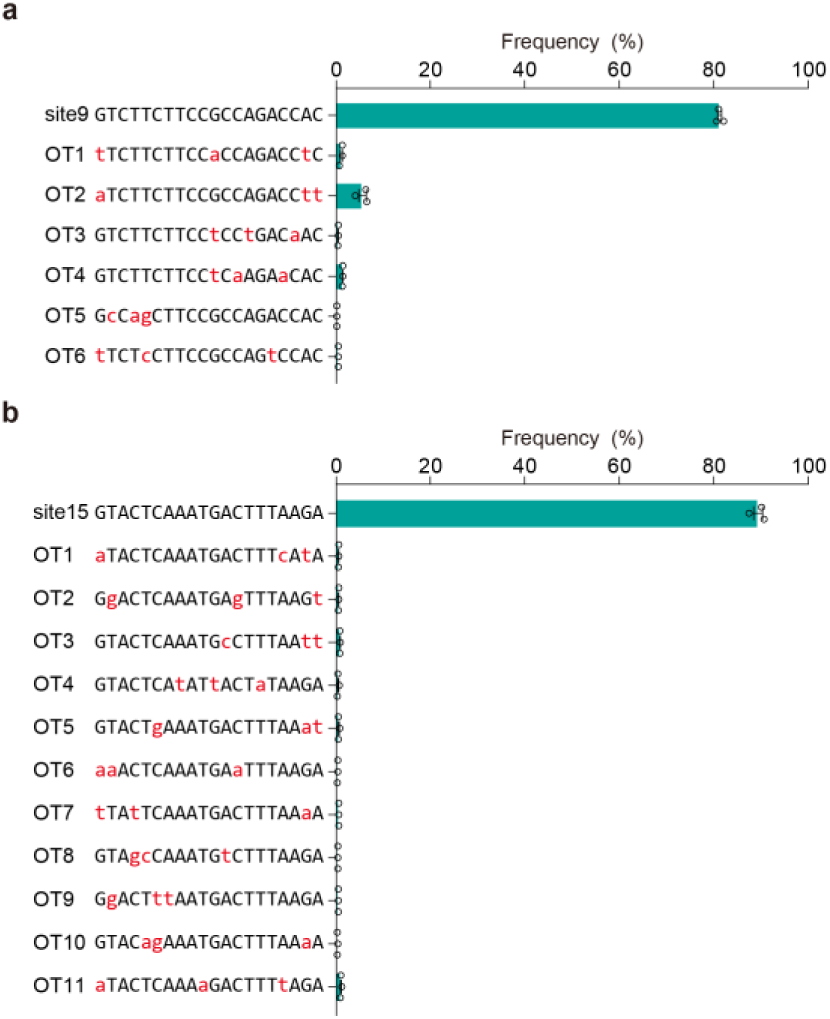
The sgRNA-dependent off-target analysis for gTBEv3 at more sites. The sgRNA-dependent off-target analysis for gTBEv3 editing at site 9 (**a**) and site 15 (**b**) (*n* = 3). OT: off-target. All values are presented as mean ±s.e.m.

**Supplementary Figure 8.**
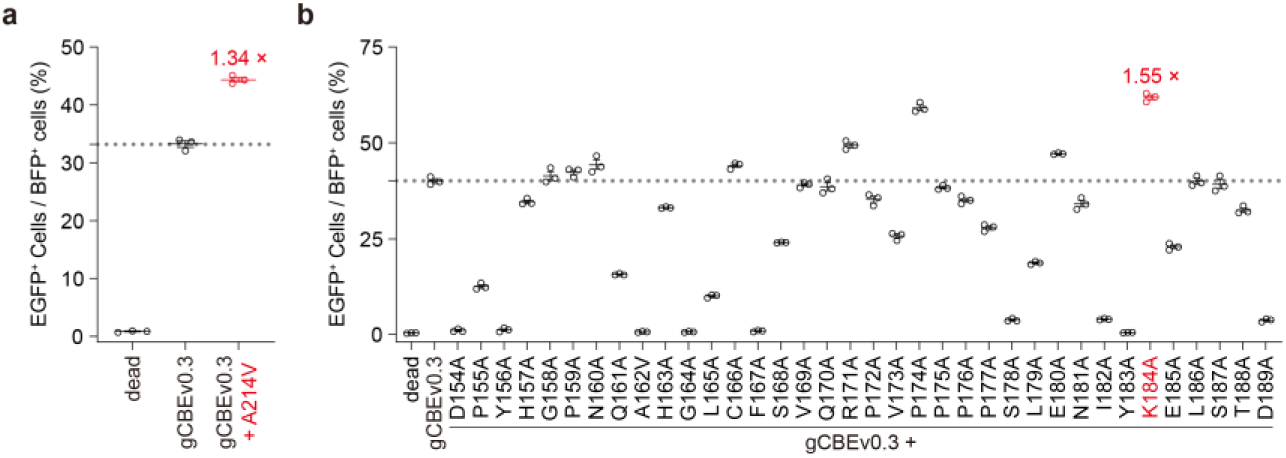
Performance of engineered variants in the gCBEv0.3 background. **a**, Percentage of EGFP^+^ cells of gCBE variant by introduction of the mutation A214V (*n* = 3). **b**, Percentage of EGFP^+^ cells of gCBE variants from alanine-scanning mutagenesis of regions covering the catalytic water-activating loop and the Pro-rich loop (*n* = 3). Replacement of alanine with valine (A>V) is intended to cover all the residues in the interested regions. dead, catalytically inactive UNG2Δ88 (carrying D154N and H277N mutations, equivalent to D145N and H268N of UNG1). All values are presented as mean ±s.e.m.

**Supplementary Figure 9.**
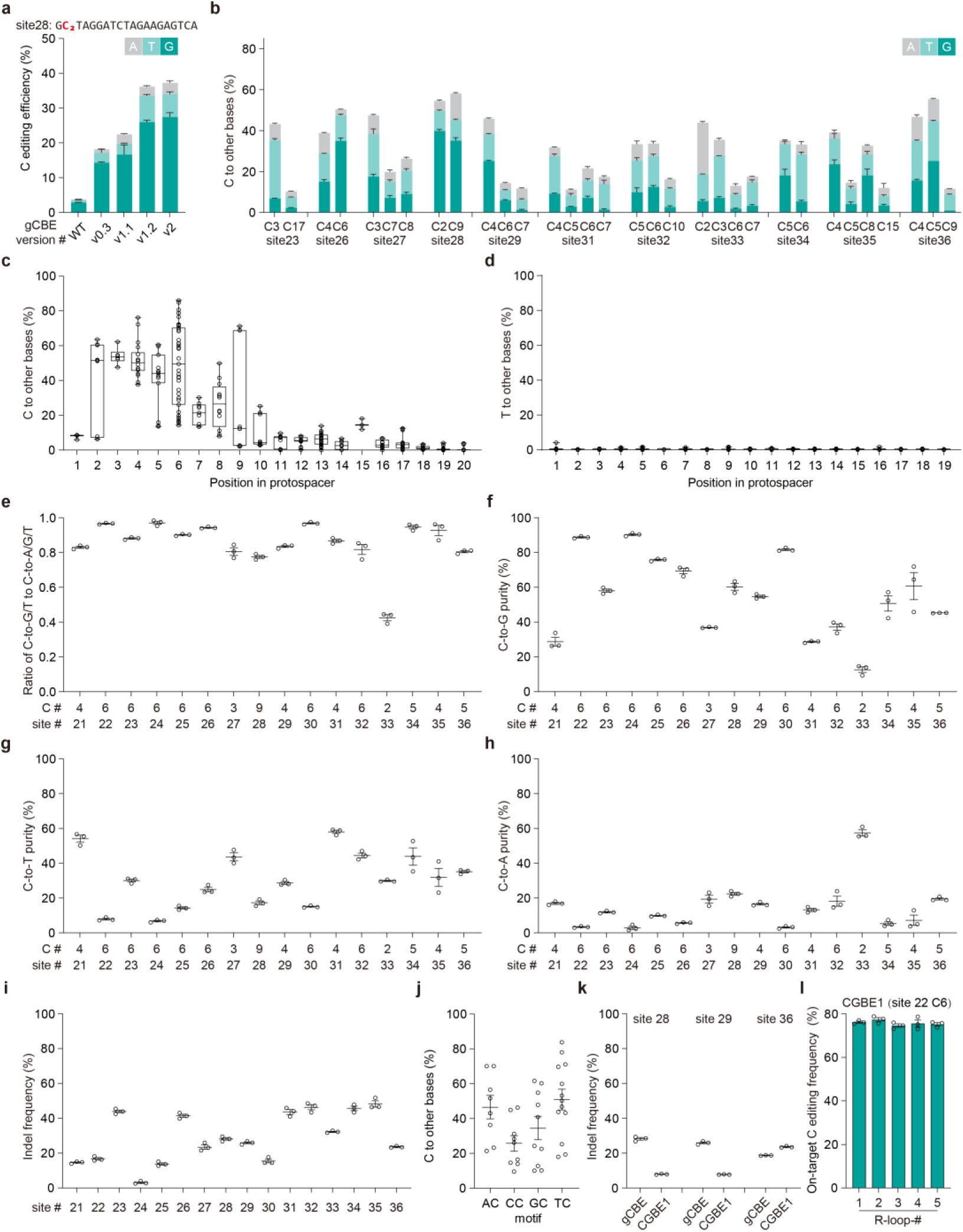
Further characterization of editing profiles for gCBEv2. **a**, Frequency of C base editing outcomes with different gCBE variants at the edited C2 position in site 28 in transfected HEK293T cells by target deep sequencing (mean ±s.e.m., *n* = 3). **b**, Bar plots showing the on-target DNA base editing at two or more positions with C conversion frequencies >10% at each genomic site in HEK293T cells (mean ±s.e.m., *n* = 3). **c,d**, Frequencies of C (**c**) and T (**d**) conversions by gCBEv2 across the protospacer positions 1-20 (where PAM is at positions 21–23) from the edited sites in Figure 4c. Single dot represents individual replicate (*n* = 3 independent replicates per site), and boxes span the interquartile range (25th to 75th percentile); horizontal lines within the boxes indicate the median (50%); and whiskers extend to the minimal and maximal values. **e**, The ratio of C-to-G/T to C-to-A/G/T conversion frequency by gCBEv2 editing at the sites shown in Figure 4c. **f-h**, Percentage of C-to-G (**f**), C-to-T (**g**) or C-to-A (**h**) editing by gCBEv2 at various edited sites shown in Figure 3a (mean ±s.e.m., *n* = 3). **i**, indels frequencies with gCBEv2 at 16 on-target sites (mean ±s.e.m., *n* = 3). In **e**-**i**, C#: C position with highest on-target base editing frequencies across protospacer positions 1-20. site #: genomic site number. **j**, The statistical analysis of on-target DNA base editing for each NC motif from the 16 edited sites. Each dot represents the mean of three biological replicates for each edited position at various edited sites. **k**, indels frequencies with gCBEv2 and CGBE1 at 3 on-target sites from Figure 4d (mean ±s.e.m., *n* = 3). **l**, On-target base editing frequencies for CGBE1 at C6 of site 22 in HEK293T cells for the orthogonal R-loop assay (mean ±s.e.m., *n* = 3).

**Supplementary Figure 10.**
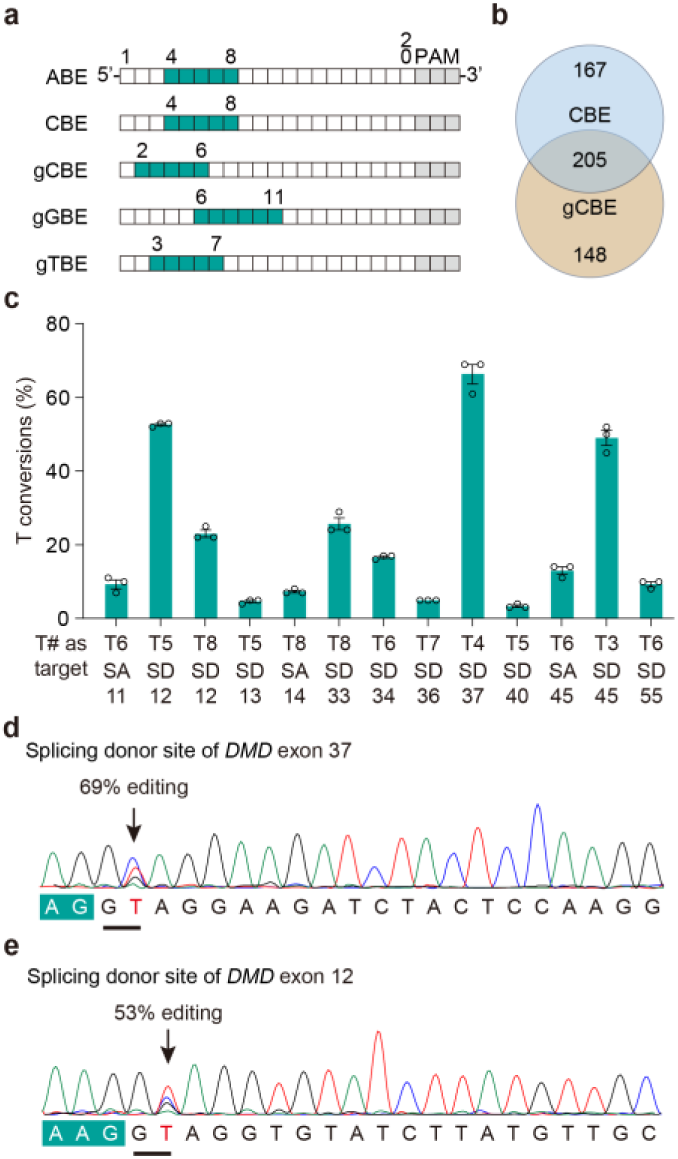
Base editing at spicing sites with gTBEv3. **a**, The optimal editing windows for various base editors. **b**, Venn diagram showing the distribution of sgRNAs for CBE and gCBE in Figure 5b. **c**, Bar plots showing the frequency of T conversations at several SA or SD sites of interest targeted by gTBEv3 (mean ±s.e.m., *n* = 3). T#: The position of targeted T across protospacer positions 1–20. Sanger sequencing results were quantified by EditR. **d**,**e**, DNA sequencing chromatograms for targeting the SD site of human *DMD* exon 37 (**d**) and exon 12 (**e**) with gTBEv3.

**Supplementary Figure 11.**
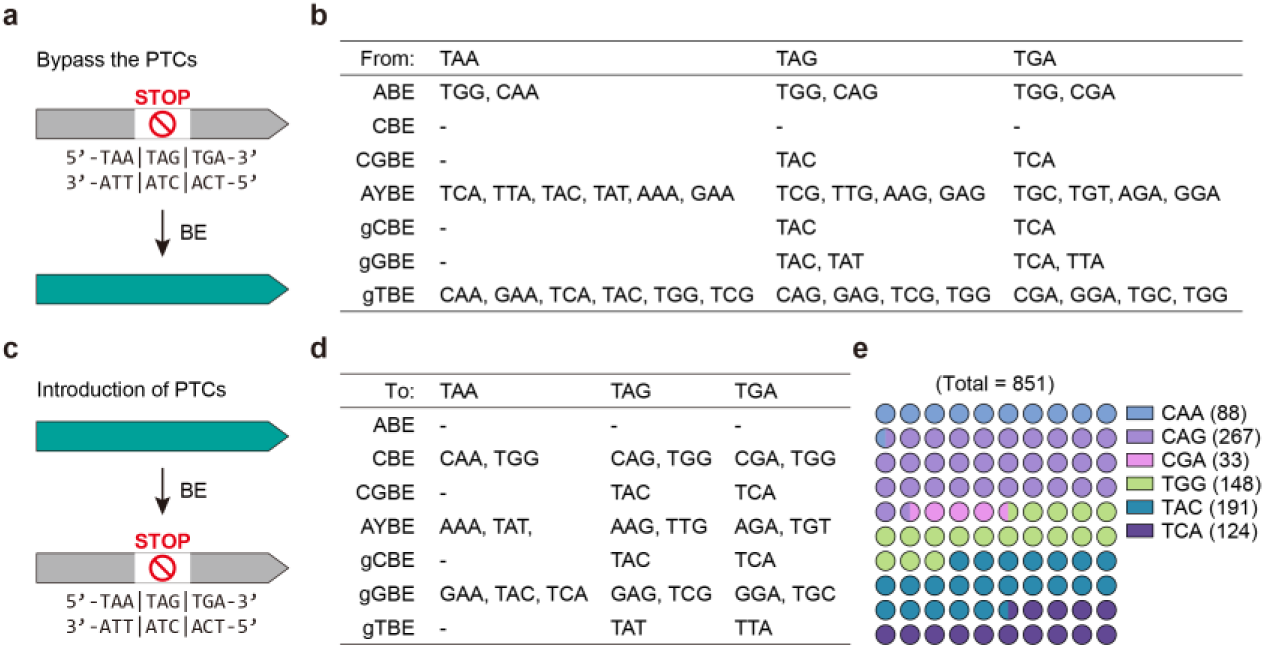
PTCs editing and introduction for various base editors. **a**, principle for bypassing premature termination codons (PTCs) with various base editors. **b**, the possible codon outcomes from stop codons (TAA, TAG or TGA) editing with different base editors. **c**, principle for introduction of PTCs with various base editors. **d**, the available codons for editing into stop codons (TAA, TAG or TGA) with different base editors. **e**, The 10 ×10 dot plot diagram showing the percentage of possible sgRNAs for introduction of premature termination codons (PTCs) by targeting different codons (with the number of available sgRNAs presented in the right) in 15 well-studied genes (*AGT, ANGPTL3, APOC3, B2M, CD33, DNMT3A, HPD, KLKB1, PCSK9, PDCD1, PRDM1, TGFBR2, TRAC, TTR, VEGFA*) for gene and cell therapy research with gGBE and CBE.

**Supplementary Figure 12.**
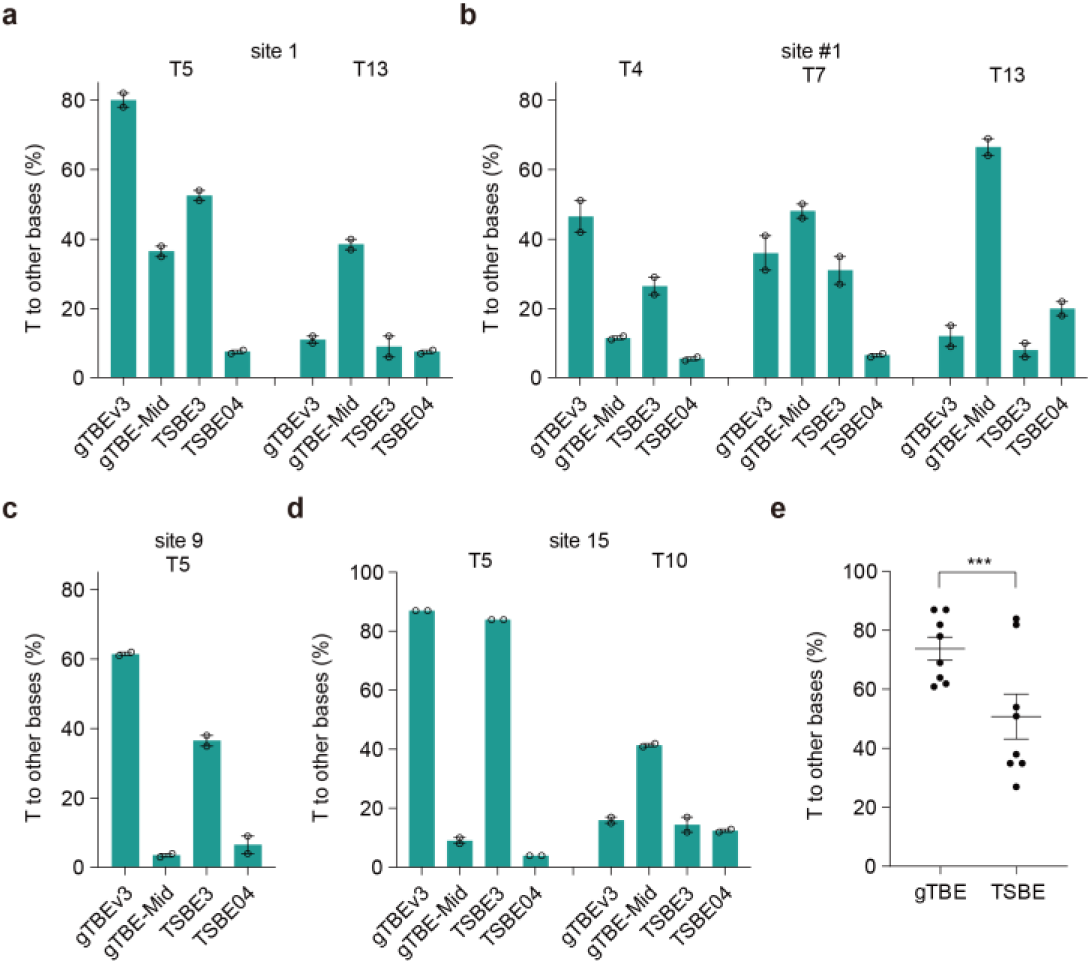
Comparison of T conversion efficiency for indicated editors. **a-d**, Bar plots showing the frequency of T conversations at T5 and T13 of site 1 (**a**), T4, T7, and T13 of site #1 (**b**), T5 of site 9 (**c**), T5 and T10 of site 15 (**d**). Single dot represents individual replicate (*n* = 2). **e,** Analysis of data from **a** to **d**. Each dot represents the individual replicate (*n* = 2) with the highest T conversion frequencies of tested sites in **a**-**d**. The gTBE represents gTBEv3 or gTBE-Mid. The TSBE represents TSBE3 or TSBE04. One-tailed paired *t*-test, ***P<0.001. gTBE-Mid represents a gTBE variant with the UNG moiety from gTBEv3 embedded into the middle of the nCas9 protein between S1248 and P1249. TSBE3 (short for TSBE3 EK) and TSBE04 (short for TSBE-CE-04) were derived from a recent preprint^43^.

